# Resilience of Atlantic Slippersnail *Crepidula fornicata* Larvae in the Face of Severe Coastal Acidification

**DOI:** 10.1101/346825

**Authors:** Nicola G. Kriefall, Jan A. Pechenik, Anthony Pires, Sarah W. Davies

## Abstract

Globally, average oceanic pH is dropping, and it will continue to decline into the foreseeable future. This ocean acidification (OA) will exacerbate the natural fluctuations in pH that nearshore ecosystems currently experience daily, potentially pushing marine organisms to their physiological limits. Adults of *Crepidula fornicata* (the Atlantic slippersnail) have proven remarkably resilient to many environmental changes, which is perhaps not surprising considering that they are common intertidally, have a geographically large native range, and have been extremely successful at invading coastal water in many other parts of the world. However, the larvae of *C. fornicata* have been shown to be somewhat more vulnerable than adults to the effects of reduced pH. Research to date has focused on the physiological impacts of OA on *C. fornicata* larvae; few studies have explored shifts in gene expression resulting from changes in pH. In the present study, we examined the response of young (4- day old) *C. fornicata* larvae to two extreme OA treatments (pH 7.5 and 7.6) relative to pH 8.0, documenting both phenotypic and genome-wide gene expression responses. We found that rearing larvae at reduced pH had subtle influences on gene expression, predominantly involving downregulation of genes related to growth and metabolism, accompanied by significantly reduced shell growth rates only for larvae reared at pH 7.5. Additionally, 10-day old larvae that had been reared at the two lower pH levels were far less likely to metamorphose within six hours when exposed to inducer. However, all larvae eventually reached similarly high levels of metamorphosis 24 hours after settlement induction. Finally, there were no observed impacts of OA on larval mortality. Taken together, our results indicate that far future OA levels have observable, but not severe, impacts on *C. fornicata* larvae, which is consistent with the resilience of this invasive snail across rapidly changing nearshore ecosystems. We propose that future work should delve further into the physiological and transcriptomic responses of all life history stages to gain a more comprehensive understanding of how OA impacts the intertidal gastropod *C. fornicata*.

## 1 Introduction

Anthropogenic combustion of fossil fuels and the subsequent rise of atmospheric partial pressure of carbon dioxide (*p*CO_2_) since the industrial revolution has introduced many threats to the marine world, including ocean acidification (OA) (Caldeira and Wickett, 2003; Bopp et al., 2013). The ocean’s average pH has already declined by 0.13 units since 1765, undergoing seawater chemistry changes unobserved for hundreds of thousands, if not millions, of years (Pelejero et al., 2010; Petit et al., 1999). Furthermore, the International Panel on Climate Change (IPCC) predicts a steeper decline of another 0.2-0.4 units by the end of this century (Plattner et al., 2001; IPCC, 2013). In estuarine and intertidal nearshore ecosystems, this anthropogenic climate change threatens to exacerbate the already substantial daily fluctuations in pH and seawater chemistry experienced in coastal waters and alter the baselines of local environmental variability (Waldbusser and Salisbury, 2014). Understanding how marine organisms will respond to these forecasted changes is a pressing conservation issue and requires a multifaceted approach including a more mechanistic understanding behind observed responses (Kroeker et al., 2013).

In response to decreased pH, many shallow water marine mollusks exhibit reduced calcification (Ries et al., 2009) and shell growth (Berge et al., 2006) rates and experience higher mortality (Shirayama and Thornton, 2005). However, compared with adult life stages, larval stages of marine organisms are typically even more vulnerable to the effects of OA (Dupont and Thorndyke, 2009; Gazeau et al., 2013; Kurihara, 2008). Marine larvae lack defenses present in adulthood (Byrne, 2011), exhibit greater surface area to mass ratios than adults, and typically require more specific environmental conditions during development (Bechmann et al., 2011; Hunt and Scheibling, 1997; Thorson, 1950), contributing to their heightened vulnerability to the impacts of OA. Negative effects of OA on larval mollusks include morphological abnormalities, reduced growth, increased mortality, and delayed metamorphosis (Dupont and Thorndyke, 2009; Gazeau et al., 2013; Kurihara, 2008). Taken together, these effects highlight the importance of studying early life stages, given that reductions in fitness in response to OA during the larval stage could have broad-reaching implications for adult populations (Dupont and Thorndyke, 2009; Gazeau et al., 2013; Kurihara, 2008).

In contrast to the more extensively studied phenotypic effects of OA on invertebrate larvae, much less in known about the genetic mechanisms underlying these responses. However, we do know that changes in gene expression in response to OA can vary widely across taxa. In sea urchin and oyster larvae, OA dampens expression of suites of genes responsible for major cellular processes, including metabolism, biomineralization, and larval shell formation (De Wit et al., 2018; Dineshram et al., 2012; O’Donnell et al., 2009; Todgham and Hofmann, 2009). Similarly, Yang et al. (2017) found significant upregulation of a tyrosinase, which is a gene important for shell repair in oyster larvae. In contrast, Zippay et al. (2010) examined two shell formation genes in red abalone larvae and determined that they were not differentially expressed at reduced pH. Furthermore, Kelly et al. (2016) found that the decreased growth rates observed for California mussel larvae in response to OA did not correlate with changes in gene expression. Interestingly, a coccolithophore was able to increase calcification over 700 generations in response to high *p*CO2 and temperature, but without differentially expressing genes traditionally associated with calcification (Benner et al., 2013).

The Atlantic slippersnail, *Crepidula fornicata*, exemplifies a marine intertidal and estuarine invertebrate whose adults exhibit remarkable resilience to most predicted environmental conditions of climate change (Diederich and Pechenik, 2013; Noisette et al., 2016; Ries et al., 2009), yet whose early life stages appear somewhat more vulnerable (Bashevkin and Pechenik, 2015; Maboloc and Chan, 2017; Noisette et al., 2014). Indeed, fecundity can be higher for intertidal individuals than for neighboring subtidal individuals (Pechenik et al., 2017b) and higher fecundities have also been observed at the northern range extreme when compared to their range center (Pechenik et al., 2017a). Adult *C. fornicata* have even demonstrated increased calcification under intermediate *p*CO_2_ conditions (605 and 903 ppm; Ries et al., 2009). Increased *p*CO_2_ (750 µatm) has also been shown to have no significant effect on *C. fornicata* rates of respiration, ammonia excretion, or filtration; indeed, a significant decrease in calcification rate was not observed until these gastropods were placed under the extremely high *p*CO_2_ of 1400 µatm (Noisette et al., 2016). Clearly *C. fornicata* adults appear remarkably robust in response to changes in pH when compared to other marine calcifying organisms (Noisette et al., 2014; Ries et al., 2009) and this resilience may help explain how *C. fornicata* has invaded diverse global environments and successfully established huge populations therein (Blanchard, 1997; Viard et al., 2006). In contrast to adult *C. fornicata*, early life stages have been shown to be relatively more negatively affected by environmental stressors. For instance, *C. fornicata* larvae under reduced pH exhibited decreased shell growth, their shells had more abnormalities, and their mineralization rates decreased (Noisette et al., 2014). A study on *C. fornicata’s* congener, *C. onyx*, found no effect of reduced pH on larval mortality, but did similarly document reduced growth rates and increasingly porous shells (Maboloc and Chan, 2017).

While researchers are beginning to characterize the morphological effects of OA on *C. fornicata* larvae, gene expression responses under reduced pH conditions remain unexplored. To better understand the response of *C. fornicata* larvae to OA, this study first characterized a suite of phenotypic traits in response to three pH treatments (pH 8.0, 7.6, 7.5) coupled with transcriptomic responses of larvae that had been reared at the different pH levels for 4 days, starting within 12 hours of hatching. Given results from previous transcriptome studies of mollusks under OA (De Wit et al., 2018; Dineshram et al., 2012), we hypothesized that marked shifts in gene expression patterns early in development would be observed, including downregulation of genes related to growth and metabolism in combination with previously reported phenotypes of reduced shell growth rates in *C. fornicata* larvae (Noisette et al., 2014). The impact of acidification on time to competence for metamorphosis was also quantified; we predicted that time to competence would be prolonged even if growth rates were unaffected, as the onset of competence relates to processes of differentiation rather than growth (Pechenik et al., 1996); delayed time to competence has previously been noted in other marine mollusks, eastern oyster and bay scallop (Talmage and Gobler, 2010). Conversely, we expected to see no impact of reduced pH on larval mortality, as has been observed in *C. onyx* larvae (Maboloc and Chan, 2017). Together, these results begin to explore the complexities of the effects of OA on an early life history stage of a highly invasive and resilient snail.

## 2 Materials and methods

### 2.1 Adult and larval collection

Several stacks of adult *C. fornicata* were collected on July 9, 2016 from the intertidal of Totten Inlet, Thurston Co., WA, and driven to the University of Washington Friday Harbor Laboratories, Friday Harbor, WA. Each stack of approximately 4-6 individuals was held unfed in a one-gallon glass jar of unfiltered seawater at room temperature (21 – 23°C); the seawater was mildly aerated and changed daily. Larvae used in this study hatched and were released naturally by a single brooding adult on the third day after adult collection. Veligers were collected by siphoning on to a 150µm sieve shortly after release.

### 2.2 Seawater pH manipulation and carbonate chemistry

Larvae were cultured in the Ocean Acidification Environmental Laboratory at the University of Washington Friday Harbor Laboratories. Incoming seawater was filtered at 1 µm and equilibrated overnight at 20°C by bubbling with ambient air (for pH 8.0) or with mixtures of CO_2_ and CO_2_-free air delivered by Aalborg GFC17 mass-flow controllers (for pH 7.5 and 7.6). Seawater pH was measured immediately before loading into culture jars with a Honeywell Durafet pH electrode calibrated to the total scale by the cresol purple spectrophotometric method described by Dickson et al. (2007). Headspaces of culture jars were continuously ventilated with the same gas mixtures used to condition the seawater pH treatments during the 2 day intervals between culture water changes. Temperature and salinity were measured with a YSI Pro Series 1030 meter. Seawater samples representing each treatment were fixed with mercuric chloride and titrated to determine total alkalinity (TA) using a Mettler DL15 automated titrator calibrated to certified reference materials (Dickson laboratory, Scripps Institution of Oceanography). *p*CO_2_ and aragonite saturation state (Ω_arg_) were estimated based on empirical measurements of pH and TA using CO_2_Sys 2.1 (Pierrot et al., 2006). Seawater chemical and physical properties used in larval culture treatments are available in Table 1.

**Table 1.** Summary of measurements (pH, salinity, and temperature) taken every 2 days per treatment, from hatching (day 0) through day 10 of the study. Total alkalinity (TA) was measured on days 2 and 10. Partial pressure of carbon dioxide (*p*CO_2_), and aragonite saturation state (Ω_arg_) were calculated from TA and pH.

| Treatment | pH | Salinity (ppt) | Temperature ($^{\circ}\text{C}$) | TA ($\mu\text{mol kg}^{-1}$) | $p\text{CO}_2$ ($\mu\text{atm}$) | $\Omega_{\text{arg}}$ |
| --- | --- | --- | --- | --- | --- | --- |
| (pH) | ± SD | ± SD | ± SD | ± SD | ± SD | ± SD |
| 7.5 | 7.53 | 29.73 | 19.93 | 2099.85 | 1480.55 | 0.83 |
| | $\pm 0.02$ | $\pm 0.39$ | $\pm 0.18$ | $\pm 3.30$ | $\pm 72.05$ | $\pm 0.04$ |
| 7.6 | 7.60 | 29.55 | 20.20 | 2083.07 | 1207.60 | 0.98 |
| | $\pm 0.01$ | $\pm 0.50$ | $\pm 0.37$ | $\pm 15.95$ | $\pm 20.43$ | $\pm 0.01$ |
| 8.0 | 7.95 | 29.53 | 20.28 | 2085.53 | 504.97 | 1.98 |
| | $\pm 0.03$ | $\pm 0.44$ | $\pm 0.12$ | $\pm 13.33$ | $\pm 35.06$ | $\pm 0.11$ |

### 2.3 Larval culture

Larvae were randomly distributed across pH treatments of 8.0, 7.6, and 7.5. Each pH treatment consisted of four replicate 800 mL jars each containing 200 larvae. Larvae were fed 15 x 10^4^ cells mL^-1^ of Tahitian *Isochrysis galbana* (T-ISO), a diet that supports maximal growth rates in larvae of *C. fornicata* (e.g. Pechenik and Tyrell, 2015). T-ISO was concentrated by centrifugation and re-suspended in filtered seawater before addition to larval cultures to minimize transfer of algal growth medium. Larvae were reared for 12 days, and sub-sampled for transcriptome analyses on day 4. Food and culture seawater were changed every 2 days. To estimate mortality, live *C. fornicata* larvae were counted during water changes on days 2, 6, and 10 and the proportion surviving on days 6 and 10 were compared to the live count taken on day 2.

### 2.4 Shell length and growth rate measurements

Growth rates of veliger larvae were measured as the change of shell length (µm) over time. Individuals were imaged using a Motic camera fitted to a Leica Wild M3C dissecting microscope, and shell lengths were measured in ImageJ. Twenty larvae were measured upon collection on the day of their hatch. After the brood was distributed into treatments, 20 larvae from each replicate culture jar were imaged and measured every 4 days until day 12 post-hatch. Larvae were blindly selected in order to avoid size biasing, and were returned to cultures after measurement.

### 2.5 Metamorphic competence measurements

When larvae were 10 days old, we exposed them to elevated levels of KCl (20 mM excess) in seawater to assess competence for metamorphosis (Pechenik et al., 2007; Pechenik and Gee, 1993; Pechenik and Heyman, 1987). For each assay, 10 larvae were taken from each of the four replicate cultures of the three pH treatments (*n* = 40 / pH treatment). The groups of 10 larvae were each pipetted into a dish (50 mL) of 20 mM excess KCl in seawater and then into another dish of the same solution (20 mL), to avoid diluting the final test solution. Percent metamorphosis was assessed 6 hours after the start of the experiment and again 24 hours later; metamorphosis was signaled by the loss of the ciliated velum.

### 2.6 Statistical analyses of phenotypic effects

All statistical tests were performed in the R environment (R Team, 2013). For larval growth, one-way ANOVAs and Tukey’s post-hoc tests were used to determine if significant differences in log-transformed cumulative growth rates (µm/day) existed between treatments at each time interval. Rates were cumulative relative to initial measurements, which were defined as day of hatching. Proportions of larvae that metamorphosed were compared via a one-way ANOVA and Tukey’s post-hoc test at 6 and 24 hours, respectively, after metamorphosis was induced in 10-day old larvae. Proportions of larvae surviving at 6 and 10 days, relative to day 2 counts were also compared via a one-way ANOVA and Tukey’s post-hoc test. All assumptions of parametric testing were explored using diagnostic plots in R (R Team, 2013).

### 2.7 RNA isolation and sequencing preparation

To achieve a sufficient quantity of RNA for whole genome gene expression profiling, three groups of 15 four-day old larvae were sampled from each of the three experimental treatments (pH 7.5, 7.6, 8.0). Total RNA was extracted from the nine samples (three treatments * three replicate groups of 15 larvae each) using RNAqueous kit (Ambion) per manufacturer’s instructions with one additional step: 0.5 mm glass beads (BioSpec) were added to the vial of lysis buffer and samples were homogenized using a bead beater for one minute. RNA quality was ascertained using gel electrophoresis by confirming the presence of ribosomal RNA bands, and 1500 ng of total RNA was used to create libraries. First, trace DNA contamination was eliminated by *DNase 1* (Ambion) digestion at 37°C for 45 min and then libraries were created as in Meyer et al. (2011) adapted for Illumina Hi-Seq sequencing (Dixon et al., 2015; Lohman et al., 2016). In brief, heat-sheared total RNA was transcribed into first-strand cDNA flanked by PCR primer sequences using oligo-dT containing primer, template-switching oligo (Matz et al., 1999) and SmartScribe reverse transcriptase (Clontech). Complementary-DNA was then PCR-amplified and Illumina barcodes were incorporated using a secondary short PCR. Samples were equalized, pooled, and size-selected prior to sequencing (single-end (SE) 50 basepair (bp)) at Tufts Genomics.

### 2.8 *Crepidula fornicata* transcriptome annotation and read mapping

Previously published *C. fornicata* contigs (Henry et al., 2010) that were >500bp in length were annotated by BLAST sequence homology searches against UniProt and Swiss-Prot NCBI NR protein databases with an *e*-value cutoff of e^−5^ and annotated sequences were assigned to Gene Ontology (GO) categories (The UniProt Consortium, 2015). Raw reads across libraries ranged from 14.2 to 25.2 million SE 50 bp sequences (Table 2). *Fastx_toolkit* was used to remove *Illumina TruSeq* adapters and poly(A)^+^ tails. Sequences <20 bp in length with <90% of bases having quality cutoff scores >20 were also trimmed. In addition, PCR duplicates were removed via a custom perl script ensuring that each read that was mapped corresponded to a unique mRNA molecule. The resulting quality filtered reads were then mapped to the *C. fornicata* transcriptome using *Bowtie2.2.0* (Langmead and Salzberg, 2012). A per-sample counts file was generated using a custom perl script that sums up reads for all genes while discarding reads mapping to more than one gene. Total *C. fornicata* mapped reads ranged from 0.5 (pH 8.0) to 1.0 (pH 7.5) million with mapping efficiencies ranging from 24.4 to 26.3% (Table 2).

**Table 2.** Summary of RNA libraries including raw single-end (SE) reads, trimmed (including de-duplication and quality filtering) reads, and mapped (to *C. fornicata* transcriptome) counts and percentages.

| Sample | # SE reads | # After<br>Trimming | # Mapped | % Mapped |
| --- | --- | --- | --- | --- |
| 7.5A | 25,224,357 | 3,222,928 | 789,550 | 24.50 |
| 7.5B | 21,750,964 | 3,591,242 | 918,366 | 25.57 |
| 7.5C | 22,895,862 | 4,078,245 | 1,027,861 | 25.20 |
| 7.6A | 23,083,355 | 3,792,669 | 926,497 | 24.43 |
| 7.6B | 16,184,588 | 3,239,532 | 816,054 | 25.19 |
| 7.6C | 17,215,957 | 3,053,984 | 770,342 | 25.22 |
| 8.0A | 15,364,186 | 2,393,789 | 612,258 | 25.58 |
| 8.0B | 19,838,321 | 2,476,114 | 651,727 | 26.32 |
| 8.0C | 14,229,958 | 1,839,873 | 480,153 | 26.10 |

### 2.9 Gene expression analysis

Differential gene expression analyses were performed with *DESeq2* v. 1.6.3 (Love et al., 2014) in R v. 3.1.1 (R Team, 2013). First, data were tested for sample outliers using the *arrayQualityMetrics* package (Kauffmann et al., 2009) and no outliers were detected. Expression data were then normalized using the *rlog* function and these normalized data (*N* = 19,717 genes) were used to broadly characterize differences in gene expression across experimental treatments using a principal component analysis with the package *vegan* (Oksanen et al., 2016). Significance was assessed using the multivariate analysis of variance function, *adonis*, in the *vegan* package (Oksanen et al., 2016). Raw count data are available in Supplementary Data Sheet 1.

*DESeq2* v. 1.6.3 was then used to identify differentially expressed genes (DEGs) consistently associated with the lower pH treatments (pH 7.5, pH 7.6) relative to pH 8.0 using generalized linear models. *DESeq2* performed automatic independent filtering to remove low abundance transcripts, which maximizes the rate of DEG discovery post multiple testing correction at an alpha of 0.1. *P*-values for significance of contrasts between the lower pH treatments (pH 7.5, pH 7.6) and pH 8.0 were generated based on Wald statistics and were adjusted for multiple testing using the false discovery rate method (Benjamini and Hochberg, 1995). Gene expression heatmaps of DEGs consistently associated with reduced pH were then generated with hierarchical clustering of expression profiles with the *pheatmap* package in R (Kolde, 2015). *DESeq2* results for both treatments are available in Supplementary Data Sheet 2 (pH 7.6) and Supplementary Data Sheet 3 (pH 7.5).

Gene ontology (GO) enrichment analysis was then performed using adaptive clustering of GO categories and Mann–Whitney U tests (GO-MWU) based on ranking of signed log *p*-values (Voolstra et al., 2011), which is particularly suitable for non-model organisms (Dixon et al., 2015). Results were plotted as dendrograms tracing the level of gene sharing between significant categories and direction of change in reduced pH treatments relative to pH 8.0 were indicated by text color.

## 3 Results

### 3.1 Rates of larval shell growth

*C. fornicata* larvae grew at a significantly slower rate in response to pH 7.5 at all three time-points after hatching when compared to larvae reared at pH 8.0, while larvae reared at pH 7.6 had a more variable response over time (Table 3; Figure 1). Specifically, on day 4, the cumulative growth rate was significantly lower for larvae reared in the pH 7.5 treatment compared to those reared at all other pH treatments (Table 3; Figure 1B). However, by day 8, larvae reared at both of the lower pH levels (7.5 & 7.6) were growing significantly more slowly than those reared at pH 8.0 (Table 3; Figure 1C). Lastly, by day 12, whereas mean shell growth rates were significantly lower for larvae reared at pH 7.5 when compared to those that had been reared at pH 7.6 and pH 8.0, there were no longer any significant differences in mean shell growth rates between larvae reared at pH 7.6 or pH 8.0 (Table 3; Figure 1D).

**Figure 1.**
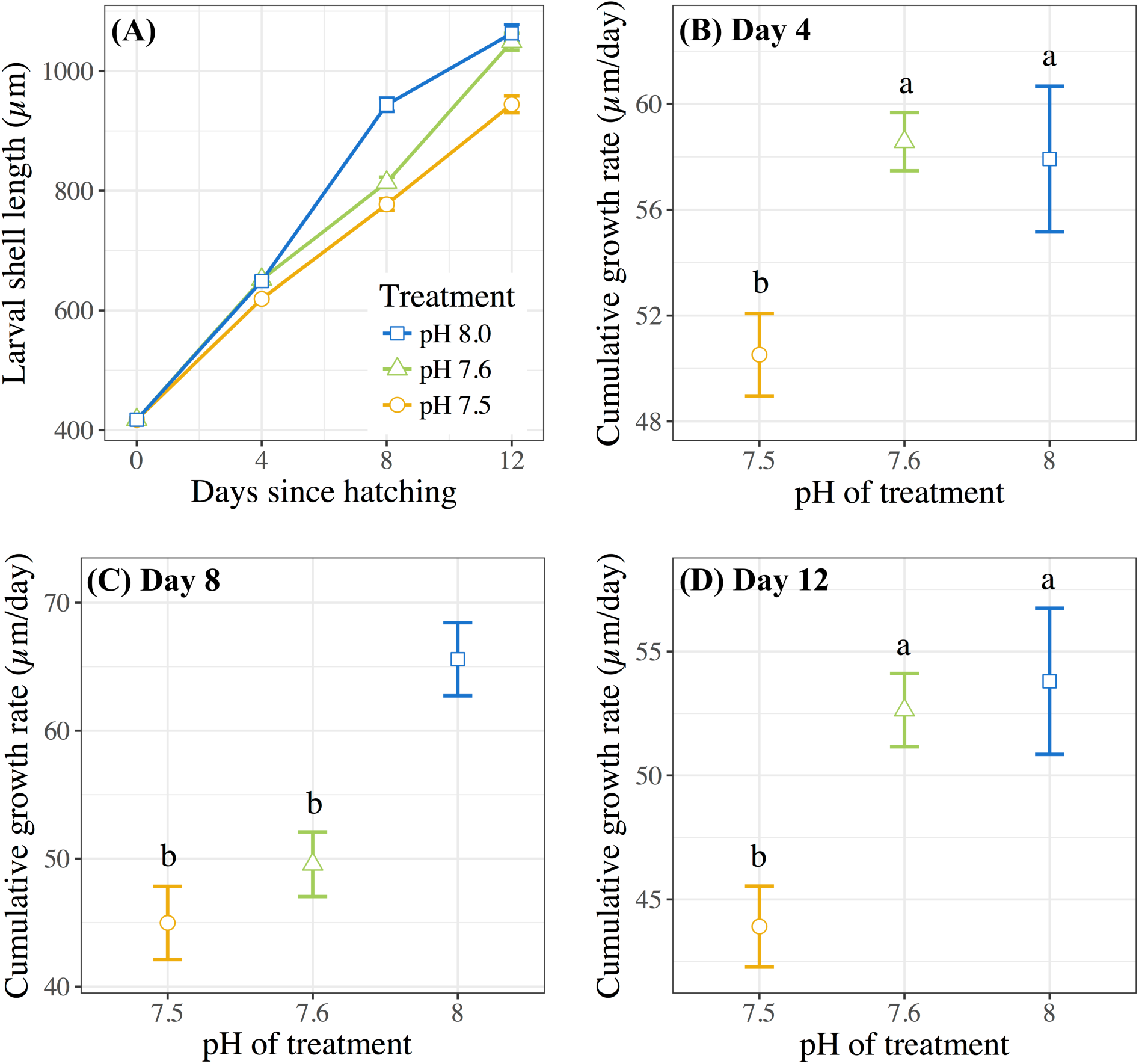
**(A)** Mean *C. fornicata* larval shell lengths (in µm) +/- SE measured 0, 4, 8, and 12 days after hatching. **(B-D)** Cumulative shell growth rates (µm/day) relative to initial (day 0) measurements compared at 4, 8, and 12 days. Different letters above bars indicate significantly different means jukey’s test; *p* < 0.05).

**Table 3.**
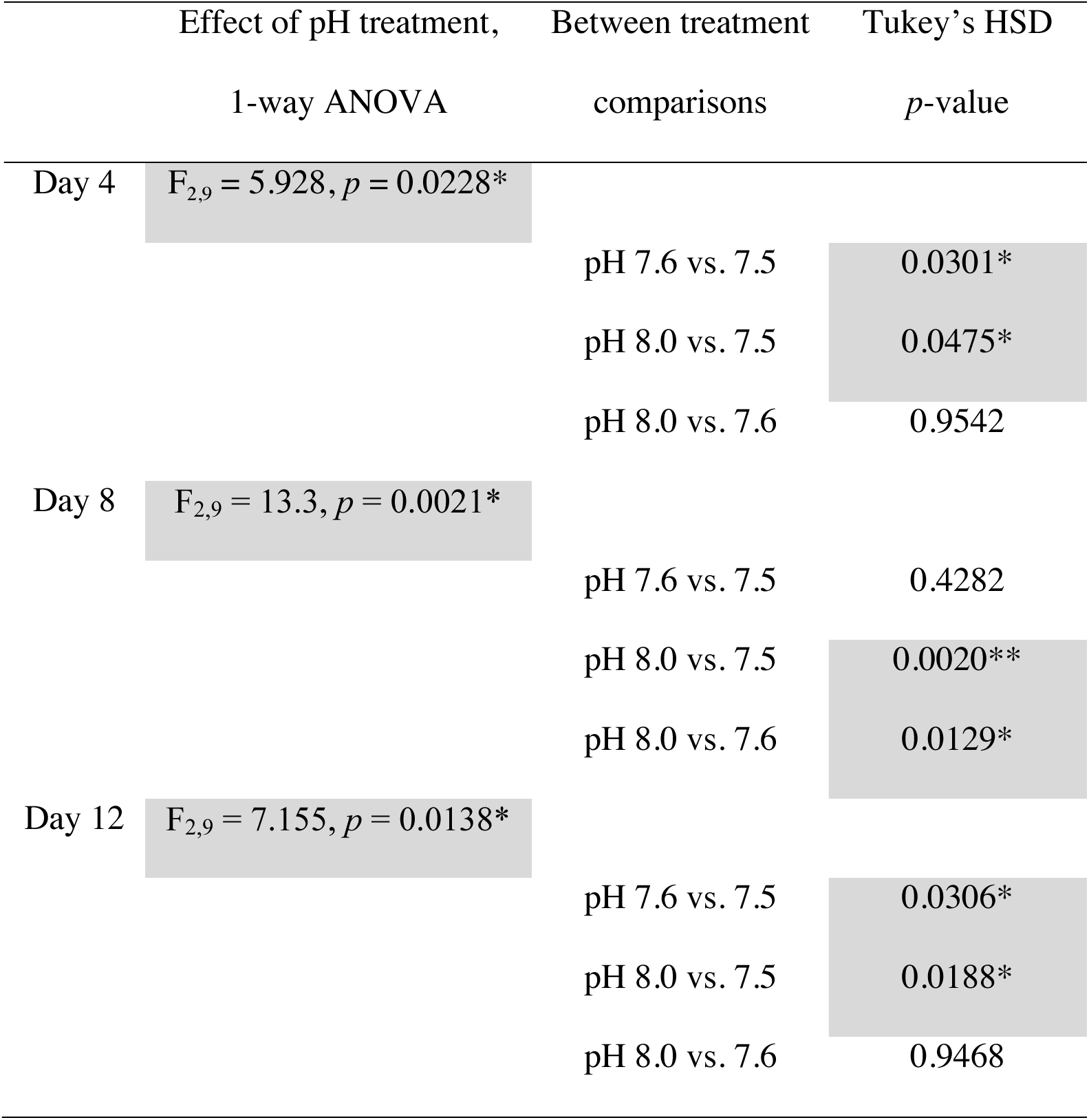
Statistical analyses of the overall effect of pH treatment and between treatment comparisons, regarding cumulative larval growth rate (µm/day) per day of measurement (days 4, 8, and 12 post-hatching). Shaded boxes indicate significant results, accompanied by the level of significance (\**p* < 0.05, \*\**p* < 0.01).

### 3.2 Metamorphic competence 10 days post-hatching

The pH in which 10-day post-hatching *C. fornicata* larvae had been reared was shown to significantly affect their ability to metamorphose within the first 6 hours of exposure to the 20 mM excess KCl inducer (Table 4; Figure 2). Larvae that had been reared at pH 7.6 or 7.5 exhibited significantly reduced metamorphosis rates at 6 hours compared to larvae reared at pH 8.0 (Table 4; Figure 2). However, 24 hours after larvae began exposure to the metamorphic inducer, larval pH treatment no longer had a significant effect on the extent of *C. fornicata* larval metamorphosis (Table 4; Figure 2).

**Figure 2.**
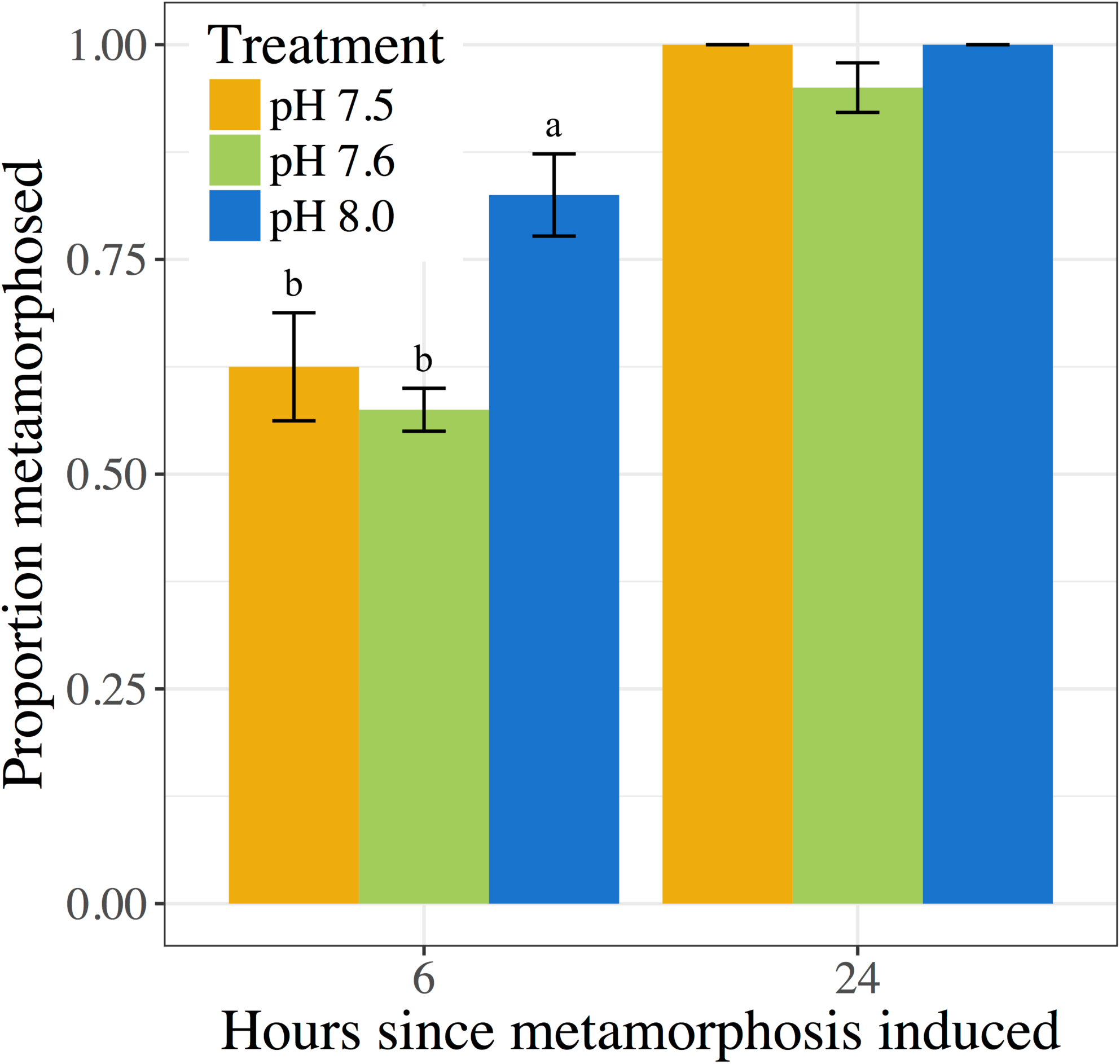
Proportion of *C. fornicata* larval metamorphosis (induced with 20 mM KCl excess) after 6 and 24 hours. Different letters above bars indicate significantly different means (Tukey’s test; *p* < 0.05). If no letters exist above bars, no significant differences were observed (Tukey’s test; *p* > 0.05).

**Table 4.** Statistical comparisons of the proportion of 10-day old larvae that metamorphosed 6 and 24 hours post-induction of metamorphosis, including overall effect of pH treatment and between treatment comparisons. Shaded boxes indicate significant results, accompanied by the level of significance (\**p* < 0.05).

| | Effect of pH treatment,<br>1-way ANOVA | Between treatment<br>comparisons | Tukey's HSD<br>$p$-value |
| --- | --- | --- | --- |
| 6 hours | $F_{2,9} = 7.636, p = 0.0115^*$ | pH 7.6 vs. 7.5 | 0.7477 |
|  |  | pH 8.0 vs. 7.5 | 0.0388* |
|  |  | pH 8.0 vs. 7.6 | 0.0124* |
| 24 hours | $F_{2,9} = 3.0, p = 0.1$ | pH 7.6 vs. 7.5 | 0.1402 |
|  |  | pH 8.0 vs. 7.5 | 1.0000 |
|  |  | pH 8.0 vs. 7.6 | 0.1402 |

### 3.3 Larval survival

Proportions of living *C. fornicata* larvae on day 6 ranged from an average of 91.6% to 97.8% across treatments (Figure 3), but there were no statistical differences in mortality rates across treatments (Table 5). The same pattern of no impact of OA on larval mortality was also observed on day 10, when proportions of living larvae ranged from 86.8% to 91.0% (Table 5, Figure 3).

**Figure 3.**
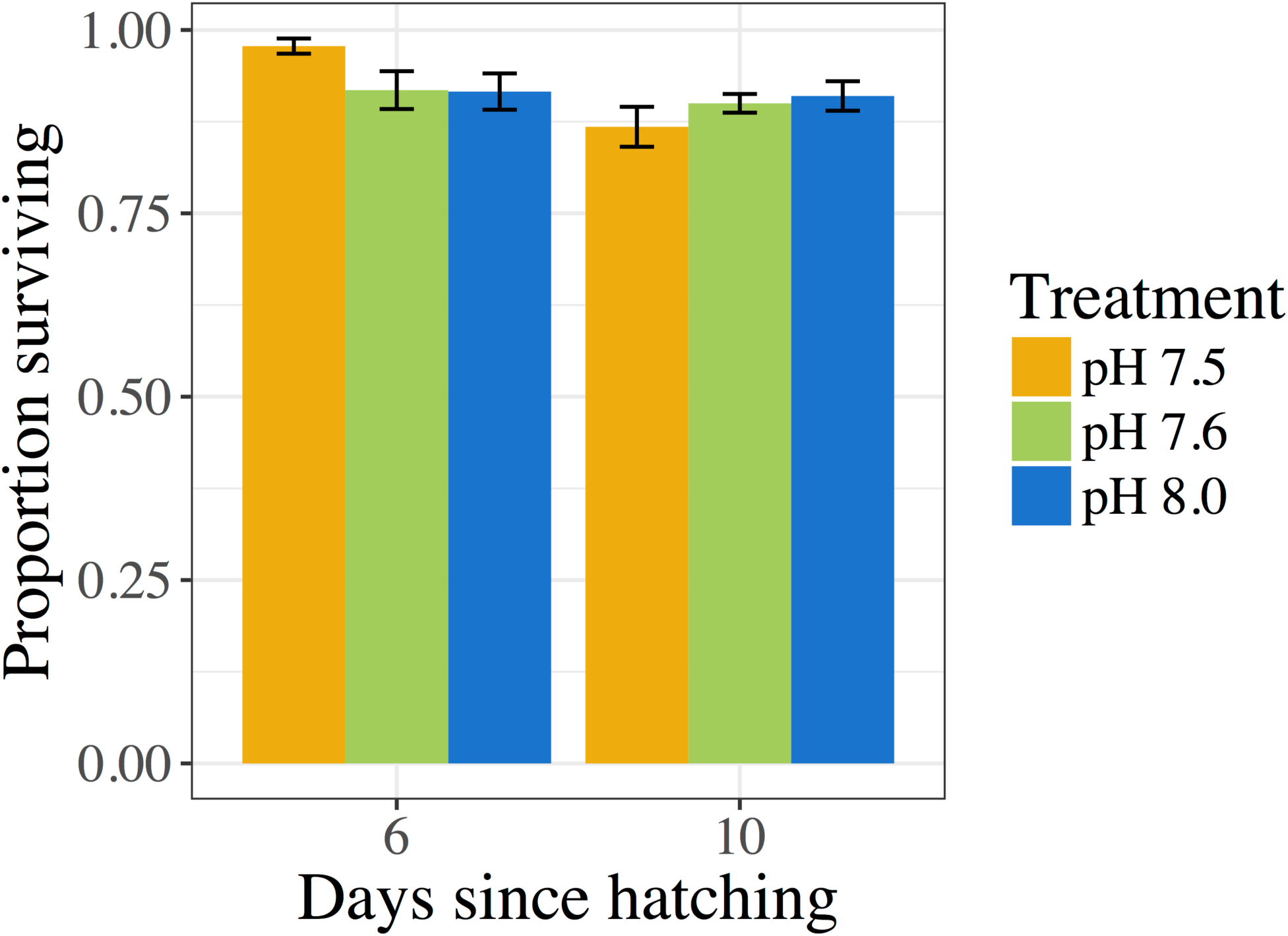
Proportion of *C. fornicata* larval survival relative to day 2, as measured on days 6 and 10. Different letters above bars indicate significantly different means jukey’s test; *p* < 0.05). If no letters exist above bars, no significant differences were observed jukey’s test; *p* > 0.05).

**Table 5.** Statistical comparisons between treatments of the cumulative proportion of surviving larvae on days 6 and 10 relative to initial counts on day 2.

|  | Effect of pH treatment,<br>1-way ANOVA | Between treatment<br>comparisons | Tukey's HSD<br>*p*-value |
| --- | --- | --- | --- |
| 6 days | $F_{2,9} = 2.736, p = 0.118$ | | |
|  |  | pH 7.6 vs. 7.5 | 0.1702 |
|  |  | pH 8.0 vs. 7.5 | 0.1537 |
|  |  | pH 8.0 vs. 7.6 | 0.9973 |
| 10 days | $F_{2,9} = 1.114, p = 0.37$ | | |
|  |  | pH 7.6 vs. 7.5 | 0.5432 |
|  |  | pH 8.0 vs. 7.5 | 0.3679 |
|  |  | pH 8.0 vs. 7.6 | 0.9395 |

### 3.4 Larval transcriptomic responses to OA

Gene expression analyses revealed strong associations between *C. fornicata* larval expression levels of transcripts within each of the three pH treatments (*Adonis p_treat_* = 0.003, Figure 4A), suggesting that each treatment elicited a distinct transcriptomic stress response. Relative to pH 8.0, *C. fornicata* larvae reared at pH 7.6 differentially expressed (FDR-adjusted <0.1) 295 genes (38 were upregulated; 257 were downregulated), while larvae reared at pH 7.5 differentially expressed 55 genes (19 were upregulated; 36 were downregulated). Notably, most differentially expressed genes (DEGs) at both pH levels (pH 7.5, 65.5%; pH 7.6, 87.1%) were underrepresented relative to pH 8.0. A total of 33 DEGs were common to larvae reared at both reduced pH levels (Figure 4B). However, rearing larvae at pH 7.6 mounted a much stronger overall transcriptomic response when compared to pH 7.5. Differential gene expression information for each treatment is included in Supplementary Data Sheets 2 and 3.

**Figure 4.**
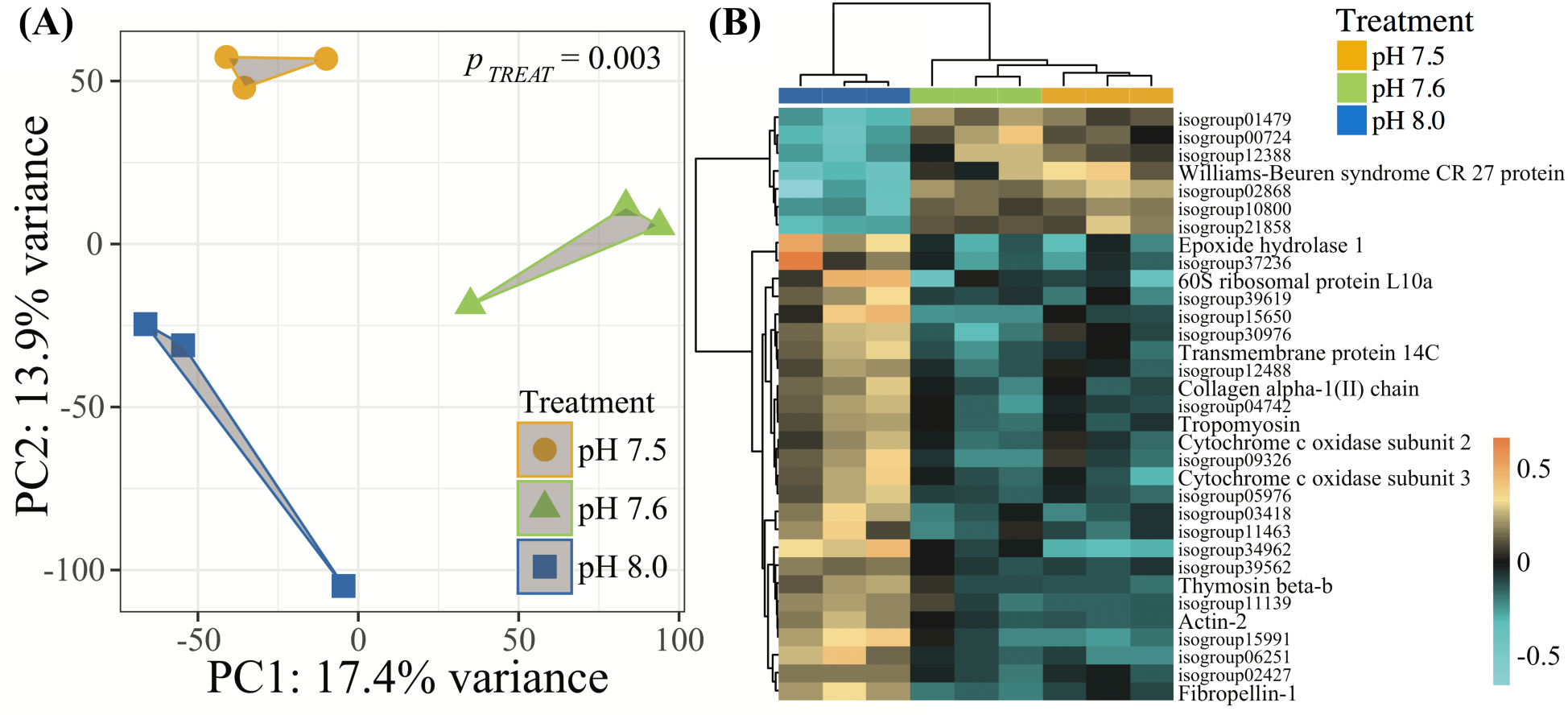
**(A)** Principle component analysis (PCA) of all r-log transformed isogroups clustered by experimental treatment, demonstrating significantly different transcriptomic responses of 4 day old *C. fornicata* larvae across different pH treatments (*Adonis p*_treat_ = 0.003). Symbol colors represent treatment conditions: blue = pH 8.0, green = pH 7.6, and yellow = pH 7.5. **(B)** Heatmap for top DEGs (FDR-adjusted = 0.10) in common for both pH 7.6 and pH 7.5 relative to pH 8.0. Rows are genes and columns are independent RNAseq libraries (N = 15 larvae/library). The color scale is in log_2_ (fold change relative to the gene’s mean). The trees are hierarchical clustering of genes and samples based on Pearson’s correlation of their expression across samples and genes.

Of the 33 DEGs in common across the two lower pH treatments, only 10 were annotated and, consistent with reduced pH causing an overall transcriptome downregulation, most of these DEGs were downregulated in larvae from reduced pH treatments relative to larvae reared at pH 8.0 (Figure 4B). These downregulated genes included genes associated with growth and metabolism (Epoxide hydrolase 1, Collagen alpha-1(II) chain, Cytochrome c oxidase subunits 2 and 3, tropomyosin, Collagen alpha-1(II) chain, Fibropellin-1), cell transport (Transmembrane protein 14C, actin-2), as well as a single gene involved in immunity (Thymosin beta-b) (Figure 4B).

### 3.5 Gene ontology (GO) enrichment in response to OA

Functional enrichments between the two reduced pH treatments (7.5, 7.6) and pH 8.0 allow for a general examination of the ‘cellular component’ (CC), ‘biological process’ (BP), and ‘molecular function’ (MF) categories that are being differentially regulated under reduced pH conditions. Consistent with *DESeq2* results, most GO enrichments were observed to be underrepresented relative to pH 8.0 (blue text; Figure 5). GO categories associated with ribosomal proteins (i.e., *ribosome;* GO:0005840, *small ribosomal subunit;* GO:0015935, *structural constituent of the ribosome;* GO:0003735), oxidative stress responses (i.e., *oxidoreductase;* GO:0016491), cytochrome-c oxidase (i.e., *cytochrome-c oxidase;* GO:0004129, *cytochrome complex;* GO:0017004), and translation activity (i.e., *translation;* GO:0006412) were consistently downregulated in *C. fornicata* larvae under lower pH treatments compared to those that had been reared under the higher pH (8.0) (Figure 5).

**Figure 5.**
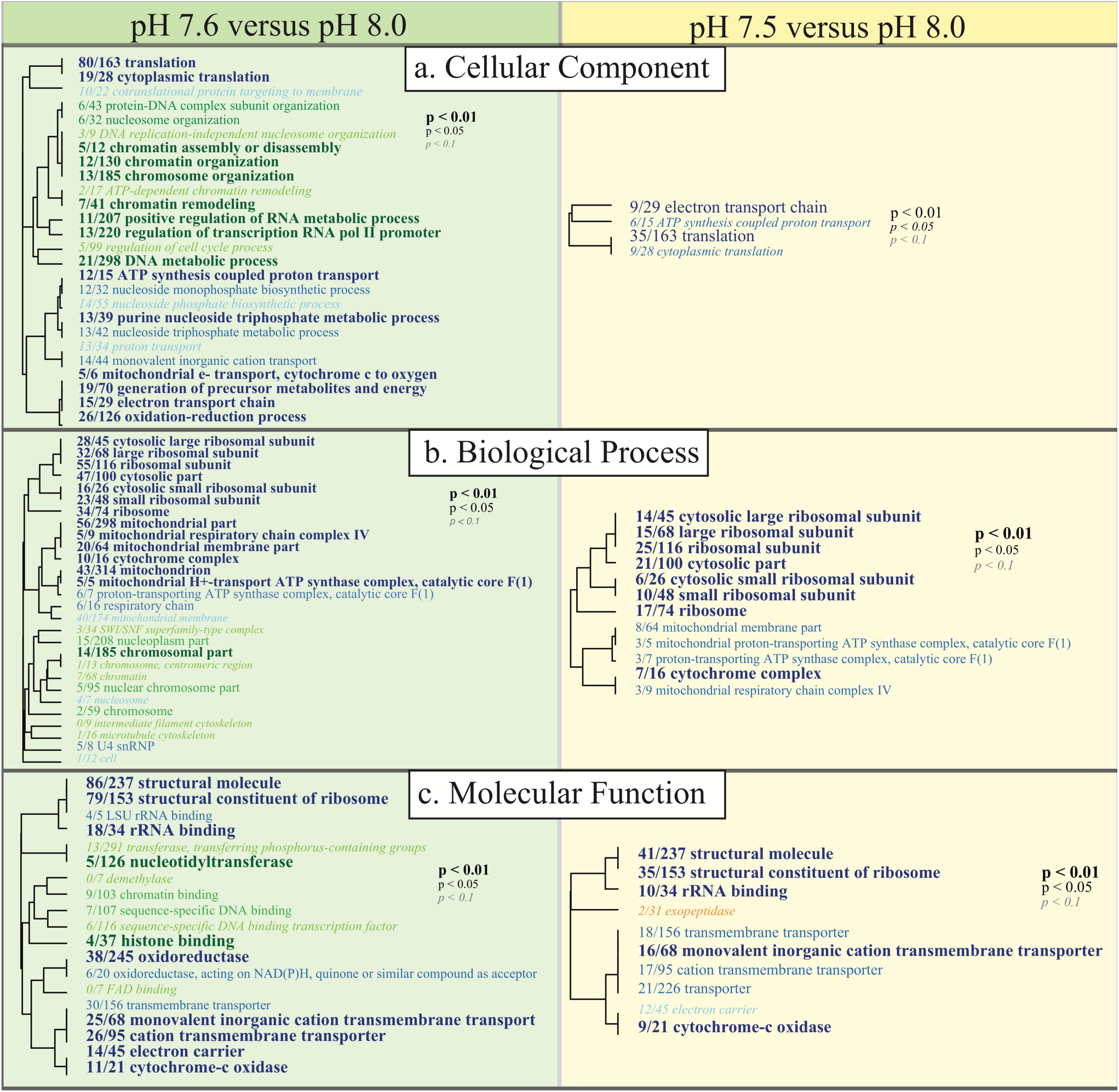
Gene ontology (GO) categories significantly enriched for each pairwise comparison (Left: pH 7.6 vs pH 8.0, Right: pH 7.5 vs pH 8.0) by **(A)** ‘cellular component’, **(B)** ‘biological process’, and **(C)** ‘molecular function’ using Mann-Whitney U tests based on ranking of signed log *p*-values. Results are plotted as dendrograms tracing the level of gene sharing between significant categories. Direction of change is noted in color where those GO categories overrepresented in pH 8.0 conditions are colored as blue and those underrepresented in pH 8.0 conditions are colored as green and yellow.

## 4 Discussion

Despite the extreme experimental levels of OA used in the present study, *C. fornicata* larvae were remarkably resilient, showing very low levels of mortality (Figure 3). Moreover, only the lowest pH treatment (pH 7.5) consistently affected larval shell growth rates (Figure 1), and short-term negative impacts of OA treatments on response to a metamorphic inducer were relieved within 24 hours (Figure 2). Moreover, surprisingly few genes exhibited differential expression amongst the experimental treatments (Figures 4, 5). Although we determined that reduced pH had a noticeable impact on some phenotypic and genotypic traits in *C. fornicata* larvae, the magnitude of the impact was relatively underwhelming. However, given that these pH conditions along with highly unsaturated aragonite are routinely experienced in Puget Sound (Bianucci et al., 2018; Reum et al., 2014), the subtle effects on physiology and transcriptomic responses perhaps might be expected for this highly resilient intertidal gastropod.

### 4.1 OA slows growth rates in *C. fornicata* larvae

Only the lowest OA treatment of pH 7.5 consistently slowed growth of *C. fornicata* larvae significantly (Figure 1), indicating the substantial extent to which the baseline environmental pH of *C. fornicata* would have to change to impact larval growth. Interestingly, larvae reared in pH 7.6 demonstrated lower growth rates between days 4 and 8 (Figure 1). It appears that, in the current study, 7.6 lies close to a threshold, below which *C. fornicata* larval growth rates were affected over the entire course of development, rather than at certain time-points (Figure 1). Accordingly, larval growth rates in the pH 7.6 treatment recovered between days 8 and 12 (Figure 1). This pattern is best explained by the Ω_arg_ concentrations hovering around 1 (aragonite saturation point) in the pH 7.6 treatment, while lying far below 1 in the pH 7.5 treatment (Table 1).

While our results indicate that larvae reared at pH 7.6 withstood effects of OA quite well, despite an early lag in growth, related studies have observed different levels of tolerance. For instance, in 14-day old *C. onyx* larvae, Maboloc & Chan (2017) found that a pH reduction to just 7.71 elicited significant decreases in shell growth rates, perhaps highlighting species-specific differences. However, complex interactions were observed between OA and diet (Maboloc and Chan, 2017). In a study exposing *C. fornicata* to OA treatments (pH 7.82, pH 7.56) over the entire course of development, from fertilization to hatching, larvae from reduced pH treatments were smaller at hatching than those under control conditions (Noisette et al., 2014), suggesting that OA impacted the reproductive or development process or that maternal effects were passed onto offspring. These two studies highlight the importance of considering other factors in addition to OA, such as pre-hatching events and diet, in future studies investigating the effects of OA on *C. fornicata.* Nevertheless, Noisette et al. (2014) conducted a literature review of the various effects of pH reductions on shell growth rates in mollusc larvae and concluded that the magnitude of diminished shell growth rates in larval *C. fornicata* was less than the observed losses across most other species, a result that corroborates the resilience to OA by *C. fornicata* larvae observed in the present study.

### 4.2 Larval *C. fornicata* transcriptomic depression in response to OA

Here we observe that after 4 days, a pH difference of just 0.1 units elicited divergent transcriptomic responses between *C. fornicata* larvae exposed to reduced pH levels (7.5 and 7.6); both responses were statistically divergent from larval expression at pH 8.0 (Figure 4). Consistent with previous research on larval responses to OA in mollusks (De Wit et al., 2018; Dineshram et al., 2012), larvae maintained at reduced pH exhibited a dampening transcriptomic response, where most differentially expressed genes were downregulated relative to pH 8.0 (Figure 4B). Amongst annotated genes that were consistently downregulated in response to OA, most were associated with growth and metabolism (Figure 4B), which corroborates findings in oyster and sea urchin larvae reared under OA conditions (Dineshram et al., 2012; Todgham and Hofmann, 2009) and is supported by the theory that metabolism cutbacks help intertidal organisms cope with short-term stress (Portner and Farrell, 2008). Gene ontology analyses also demonstrated a pattern of transcriptomic depression, where most of the significantly enriched GO terms associated with metabolism and growth (e.g. ribosomal and mitochondrial functions, ATP-synthase activity) were downregulated relative to pH 8.0 (Figure 5).

Downregulation of genes associated with growth and metabolism is also consistent with our phenotypic results in terms of shell growth rate depression in larvae reared at pH 7.5 (Figure 1). However, on day 4 (the day of transcriptomic sampling), larvae reared at pH 7.6 showed more differential gene expression than larvae reared at pH 7.5, but without the same observed repression of shell growth (Figures 1, 5). It is possible that the significant changes in gene expression on day 4 in the pH 7.6 treatment preclude the cumulative reduction in shell growth rates observed on day 8 of this experiment (Figure 1). Alternatively, pH 7.6 could represent a pH in which larvae were inducing many physiological changes to compensate for the pH treatment, which would be consistent with the observed fluctuations in shell growth rates observed at pH 7.6 and in the larger number of differentially enriched GO terms (Figure 5). Future work should focus on sampling *C. fornicata* larvae with increased temporal frequency to discern with more acuity the timing at which the strongest coinciding phenotypic and transcriptomic impacts are mounted in response to OA.

Additionally, *C. fornicata* larvae reared in the pH 7.6 treatment downregulated redox-regulation associated genes (see oxidoreductase and oxidation-reduction process in Figure 5), which are typically associated with the minimal cellular stress response (Kültz, 2005). It appears contradictory that marine organisms would downregulate these genes while under pH stress; indeed, upregulation rather than downregulation of redox functional processes has been reported in Emerald rockcod *Trematomus bernacchii* (Huth and Place, 2016) and in marine copepods (Lauritano et al., 2012) exposed to reduced pH. However, downregulation of stress response genes under OA conditions is shared among many other marine invertebrates, including purple sea urchin *Strongylocentrotus purpuratus* larvae (Todgham and Hofmann, 2009), *Saccostrea glomerata* oysters (Goncalves et al., 2017), Arctic copepods *Calanus glacialis* (Bailey et al., 2017), and coral *Acropora millepora* (Kaniewska et al., 2012). Downregulation of stress response genes may leave cells more vulnerable to the negative impacts of oxidative stress and protein denaturing (Todgham and Hofmann, 2009). Unexpectedly however, downregulation of stress response genes in response to OA did not present with any associated phenotypic impacts in *C. glacialis* copepods (Bailey et al., 2017), has been associated with pH-resistant oysters (Goncalves et al., 2017), and the *C. fornicata* larvae reared in the pH 7.6 treatment in the present study did not present with atypical growth rates at the time of transcriptome sampling, confirming that some pH-tolerant organisms typically downregulate redox-associated genes (Bailey et al., 2017).

### 4.3 OA delays larval metamorphosis

*C. fornicata* larvae, if competent, typically metamorphose within 6 hours when exposed to seawater with KCl concentrations elevated by 15 – 20 mM (Pechenik and Gee, 1993; Pechenik and Heyman, 1987). In our study, a smaller percentage of larvae that had been reared at reduced pH metamorphosed in response to KCl within 6 hours, but there was no significant effect of rearing conditions on percent metamorphosis by 24 hours (Figure 2). The reduced response to excess KCl in the first 6 hours for larvae reared at reduced pH could reflect a slightly longer time to become competent to metamorphose or interference with the functioning of the metamorphic pathway. This issue requires further study.

Similar and more extended delays in metamorphosis have been observed in other marine calcifiers, including hard clams, bay scallops, and coral larvae (Nakamura et al., 2011; Talmage and Gobler, 2010). In coral *Acropora digitifera* larvae, pH levels of 7.6 and 7.3 significantly reduced metamorphosis after both 2 hours and 7 days of OA treatment, indicating that metamorphosis may be fully disrupted or severely delayed in these larvae (Nakamura et al., 2011). In the hard clam *Merceneria merceneria*, less than 7% of larvae metamorphosed after 14 days of development at pH 8.05, 7.80, and 7.53, compared to 51% metamorphosis in pH 8.17 (pre-industrial conditions) (Talmage and Gobler, 2010). After 12 days of development, 87% of bay scallop *Argopecten irradians* larvae metamorphosed in pre-industrial pH treatments as compared to 68% metamorphosis in pH 8.05 (Talmage and Gobler, 2010). Metamorphosis percentages in both species appeared approximately the same in all treatments by day 30 for *M. merceneria* and by day 19 for *A. irradians* (Talmage and Gobler, 2010). However, the current study did not examine percentage of metamorphosis while in natural seawater conditions, nor in the natural inducer of adult-conditioned seawater (Pechenik and Heyman, 1987; Pechenik and Gee, 1993).

While a slight temporal delay in metamorphosis of *C. fornicata* larvae may seem inconsequential, it is theorized that additional time spent in the water column will increase predation rates and size-specific mortality or could carry larvae away from suitable habitat prior to metamorphosis (Bashevkin and Pechenik, 2015; Pechenik, 1999; Talmage and Gobler, 2010). On the other hand, currents could carry larvae to more suitable habitat and further aid in the invasive dispersal potential of *C. fornicata* (Bashevkin and Pechenik, 2015; Pechenik, 1999). Future work should examine the interplay of the impacts of OA with the dynamics of larval settlement and predation rates.

### 4.4 Conclusions

Higher resistance to the negative impacts of environmental stressors when compared to native species is one of the defining characteristics of a successful invasive species (Lenz et al., 2011). Accordingly, in the presence of severe OA stress, *C. fornicata* larvae in the present study exhibited some deleterious but overall moderate effects, especially when compared with effects previously documented for other marine invertebrates. As oceans become more acidic, it is likely that *C. fornicata* populations, which are already able to withstand many extreme environmental conditions today (Diederich and Pechenik, 2013; Noisette et al., 2016) may be better able to outcompete native species. To actualize a more complete picture of how OA impacts *C. fornicata* early life stages, future work should more thoroughly characterize how quickly shifts in gene expression take place in response to OA, examine more diverse life stages, and consider the potential additional impacts of climate change (temperature and nutrient shifts). While the present study only considered the impact of reduced pH, Pechenik (1984) and Bashevkin and Pechenik (2015) found that warming temperatures favored *C. fornicata* larval growth, while Thieltges et al. (2004) suggested that colder temperatures have previously prevented range expansion of *C. fornicata* into northern Europe. Considering these results in conjunction with the present study, large populations of this resilient snail may be supported well into the future and perhaps even into new areas of the globe.

## 6 Author contributions statement

AP and JP conceptualized the experimental design and carried out the organismal collections and phenotypic measurements. Transcriptomic data collection and analyses were performed by SD. Phenotypic data analyses were performed by NK. NK wrote the manuscript with contributions from SD, AP, and JP. All authors revised and approved the manuscript.

## 8 Acknowledgements

We thank Christine Choi, Rulaiha Taylor, and Alissa Resnikoff for assisting with maintenance of the larval cultures at Friday Harbor Laboratories. We also acknowledge the Marchetti and Castillo labs at UNC Chapel Hill for use of their lab spaces for the preparations of TagSeq libraries.

## 9 Data availability statement

Detailed protocol of library preparation and associated bioinformatics can be found at https://github.com/z0on/tag-based_RNAseq. Protocol of gene ontology analysis can be found at https://github.com/z0on/GO_MWU. All annotation files for *Crepidula fornicata* transcriptome are available: http://sites.bu.edu/davieslab/data-code. Raw reads have been submitted to SRA under PRJNA471890. All phenotypic and seawater measurement datasets are available upon request.

